# Structural insights into the *in situ* assembly of clustered protocadherin γB4

**DOI:** 10.1101/2024.07.05.602218

**Authors:** Ze Zhang, Fabao Chen, Zihan Zhang, Luqiang Guo, Tingting Feng, Zhen Fang, Lihui Xin, Yang Yu, Hongyu Hu, Yongning He

## Abstract

Clustered protocadherins (cPcdhs) belong to the cadherin superfamily and play important roles in neural development. cPcdhs can mediate homophilic adhesion and lead to self-avoidance and tiling by giving neurons specific identities in vertebrates. Structures and functions of cPcdhs have been studied extensively in the past decades, but the mechanisms behind the functions have not been fully understood. Here we investigate the *in situ* assembly of cPcdh-γB4, a member in the γ subfamily of cPcdhs, by electron tomography and find that the full length cPcdh-γB4 does not show regular organization at the adhesion interfaces. By contrast, cPcdh-γB4 lacking the intracellular domain can generate an ordered zigzag pattern between cells and the *cis* interacting mode is different from the crystal packing of the ectodomain. We also identify the residues on the ectodomain that might be important for the zigzag pattern formation by mutagenesis. Furthermore, truncation mutants of the intracellular domain of cPcdh-γB4 reveal different assembly patterns between cell membranes, suggesting that the intracellular domain plays a crucial role in the intermembrane organization of cPcdh-γB4. Taken together, these results suggest both ectodomain and intracellular domain regulate the *in situ* assembly of cPcdh-γB4 at the adhesion interfaces, thereby providing mechanistic insights into the functional roles of cPcdhs during neuronal wiring.

## Introduction

During neural development, neurons are organized into complex networks by following certain repulsive interactions, including self-avoidance and tiling, to guarantee correct arrangements and functionality of the networks (Lin et al., 2004; Sagasti et al., 2005; Sugimura et al., 2003). Self-avoidance refers to the repulsion between arbors from a single neuron, during which neurons need to discriminate self from non-self (Kramer and Kuwada, 1983; Kramer and Stent, 1985). Therefore, self-avoidance demands that a single neuron has its own specific identity distinct from thousands of others it may contact (Mountoufaris et al., 2018; Zipursky and Sanes, 2010). In tiling, different neurons with the same functional roles would avoid each other by sharing the same identities (Chen et al., 2017b; Grueber and Sagasti, 2010).

In *Drosophila,* Down syndrome cell adhesion molecules 1 (DSCAM1) and DSCAM2 have been shown to play key roles in self-avoidance (Hattori et al., 2009; Hattori et al., 2007; Matthews et al., 2007) and tiling (Millard et al., 2007), respectively. In vertebrates, evidence suggests that self-avoidance and tiling are mediated by cPcdhs, which can lead to repulsion between axonal or dendritic neurites (Chen et al., 2017b; Ing-Esteves et al., 2018; Kostadinov and Sanes, 2015; Lefebvre et al., 2012; Mountoufaris et al., 2017). cPcdhs belong to the cadherin superfamily and are named according to the clustered genomic organization (Wu and Maniatis, 1999) and general existence in distantly related species (Sano et al., 1993). cPcdhs contain 50-60 isoforms, and the genes of cPcdhs locate on human chromosome 5 (Wu and Maniatis, 1999) or mouse chromosome 18 (Wu et al., 2001) and are arranged closely in three tandem clusters, which correspond to three subfamilies: α, β, and γ. Each cluster contains 10∼30 variable exons and each variable exon encodes an intact ectodomain (EC), a transmembrane domain (TM) and a variable intracellular domain (VIC). Variable exons in γ cluster can be furtherly divided into type A and type B (Wu and Maniatis, 1999; Wu et al., 2001). α and γ clusters also contain three constant exons, which encode a common intracellular domain (CIC) that is conserved in all isoforms within the cPcdh subfamilies (Mah and Weiner, 2017; Mountoufaris et al., 2018). Therefore the intracellular domains (IC) of α and γ-cPcdhs have both VIC and CIC, while β-cPcdhs lack the cluster-specific CIC (Pancho et al., 2020).

To achieve self-avoidance and tiling, cell adhesion mediated by the adhesion molecules is required for both processes (Millard et al., 2007; Mountoufaris et al., 2018; Wu et al., 2012). Published data have shown that cells expressing the same sets of cPcdh isoforms exhibit cell adhesion, while a single isoform mismatch in the combinations would abolish adhesion (Thu et al., 2014). Such high matching demand between repertoires means that 50∼60 cPcdh isoforms can support identities for billions of neurons (Honig and Shapiro, 2020; Rubinstein et al., 2015; Zipursky and Grueber, 2013).

The homophilic binding of cPcdhs has been studied extensively in the past decades (Goodman et al., 2022; Goodman et al., 2016a; Rubinstein et al., 2015; Schreiner and Weiner, 2010; Thu et al., 2014). The ectodomains of cPcdhs contain six extracellular cadherin domains (EC1 to 6). Crystallographic results show that the *trans* homophilic interaction occurs between EC1-4 of the monomers (Goodman et al., 2022; Goodman et al., 2016a; Goodman et al., 2016b), while EC5-6 may mediate the *cis*-dimer formation (Goodman et al., 2022; Goodman et al., 2017; Goodman et al., 2016b; Rubinstein et al., 2015). The alternate *trans* and *cis* interactions of cPcdh ectodomains may result in an extended zipper-like structure between membranes as has been shown in a liposome model (Brasch et al., 2019). Imaging characterizations of the cells expressing cPcdhs have also been reported before (Hanson et al., 2010), but details at the adhesion interfaces remain unclear.

In the meantime, evidence has shown that the intracellular domains of cPcdhs are involved in the activation of downstream signaling cascades (Chen et al., 2009; Keeler et al., 2015; Li et al., 2012; Lin et al., 2010; Mah et al., 2016), which could be important for neuronal avoidance (Honig and Shapiro, 2020; Pancho et al., 2020).

Moreover, biochemical assays suggest that the intracellular domains of cPcdhs could interact with each other (Murata et al., 2004; O’Leary et al., 2011; Shonubi et al., 2015) and may also restrict accumulation of cPcdhs at cell-cell contacts (Fernandez-Monreal et al., 2009; Schreiner and Weiner, 2010), but the exact roles of intracellular domain in adhesion or self-avoidance have not been fully understood.

Here we explore the *in situ* assembly of cPcdh-γB4 (γB4) by combining fluorescence microscopy, electron tomography (ET) and mutagenesis studies, which would provide insights for the mechanism of γB4 in mediating cell adhesion and neuronal avoidance during neural network formation.

## Results

### Full length γB4 does not form an ordered assembly pattern at the adhesion interfaces

Crystal structure shows that the ectodomain of γB4 can generate a zipper-like pattern through alternate *trans*-interaction of EC1-4 and *cis*-interaction of EC5-6, this assembly feature was also observed in a liposome modeling system where the ectodomain of γB6 was coupled onto the liposome surfaces (Brasch et al., 2019). In order to examine the *in situ* assembly of γB4, we transfected HEK293 cells with the full length mouse γB4 (γB4-FL) fused with a GFP tag at the C-terminus, and fluorescent confocal microscopy was applied to monitor the formation of cell adhesion. Images showed that green fluorescent lines were highlighted at cell-cell contacts where γB4 accumulated for adhesion (Fig. 1A, S1A). Then the transfected cells were subjected to high-pressure freezing and freeze substitution (HPF-FS), and the plastic embedded ultra-thin sections were prepared for electron microscopic (EM) observation (Chang et al., 2018; Guo et al., 2021; He and He, 2014). The resulting EM images displayed some electron-dense features between the adjacent cell membranes at cell-cell contact regions (Fig. 1A, S1B, C), which was not observed for the non-transfected cells (Guo et al., 2021). However, no ordered assembly pattern was found after inspecting a number of adhesion interfaces (more than 10 interfaces). Since the EM sections were prepared by random cuts in 3D, we also checked the interfaces with different viewing angles by rotating the specimens in electron microscope, and no regular pattern was observed in the interfaces (Fig. 1B).

**Figure 1.**
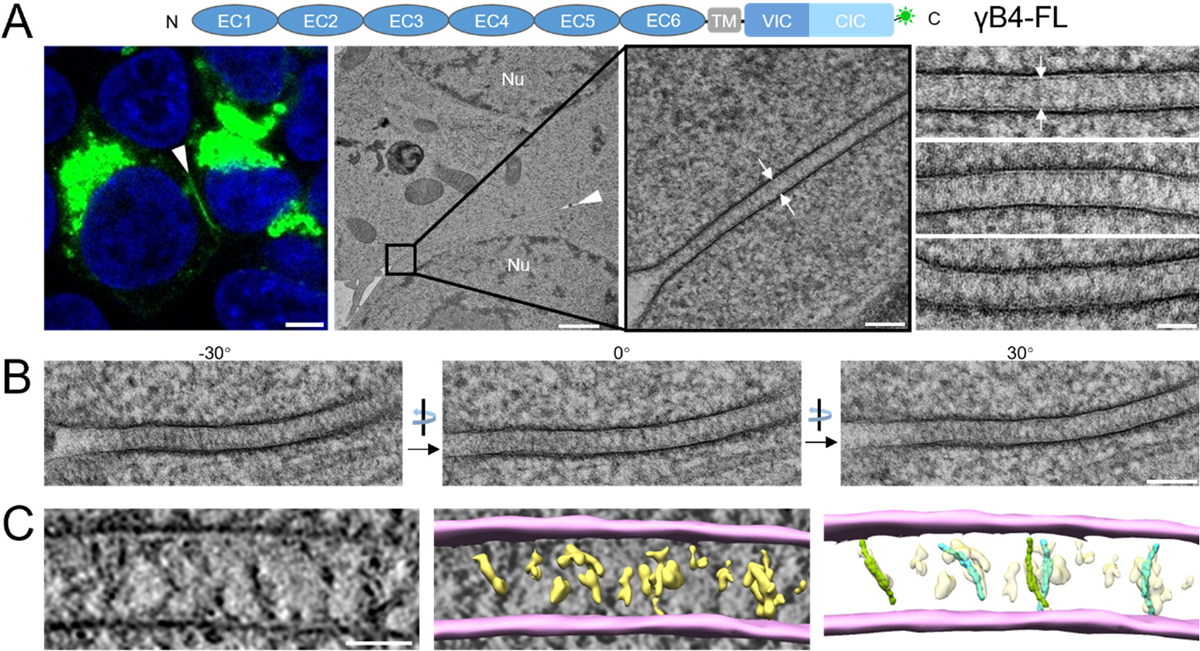
Microscopic images of the cell adhesion interfaces by γB4-FL. (A) A schematic diagram of the domain arrangement of γB4-FL is shown on the top, the GFP tag is shown in green. A confocal fluorescent image of an adhesion interface (white arrowhead) by γB4-FL is shown on the left (scale bar, 5 μm). EM images of an adhesion interface (white arrowhead) (scale bar, 1 μm) with a zoom-in view (white arrows) (scale bar, 100 nm) are shown in the middle. A gallery of the γB4-FL mediated adhesion interfaces (white arrows) is shown on the right (scale bar, 50 nm). (B) EM images of a γB4-FL mediated adhesion interface visualized at different tilt angles (scale bar, 100 nm). (C) A tomographic slice of a γB4-FL mediated adhesion interface (left) (scale bar, 35 nm) and a segmentation model of the tomogram (middle). The cell membranes and the densities in between are colored in pink and yellow, respectively. The densities are tentatively docked with the *trans*-dimers of the ectodomain of γB4 (green or cyan) (right).

Furthermore, EM tilt series were collected for tomographic reconstruction, the resulting tomograms confirmed that no ordered structure was assembled at the adhesion interfaces (Fig. 1C). After semi-automated segmentation of the tomograms (Chen et al., 2017a), a few density volumes that may correspond to the *trans*-dimers of the ectodomain of γB4 could be observed and were tentatively docked by the crystal structure with poor accuracy (Fig. 1C). These data suggest that the *in situ* organization of γB4-FL at the adhesion interfaces might be different from the crystal packing. In addition, the tomograms showed that the intermembrane distance of the adhesion interfaces mediated by γB4-FL was about 34 nm (Fig.7E), rather than 38 nm according to the assembly model based on the crystal packing of γB4 ectodomain (Fig. 8A).

### γB4 lacking the intracellular domain forms an ordered zigzag pattern at the adhesion interfaces

In parallel with the experiments for γB4-FL, we also transfected HEK293 cells with the γB4 lacking the intracellular domain (γB4-ΔIC) and prepared the specimens similarly. Fluorescent confocal images confirmed that γB4-ΔIC also accumulated at the adhesion interfaces, forming highlighted green lines (Fig. 2A, S2A). But surprisingly, EM images showed that γB4-ΔIC formed an ordered zigzag pattern between cell membranes at the adhesion interfaces (Fig. 2A, S2B, C). And we also found that the patterns could vary for different interfaces, which might be due to the different cutting angles during EM sectioning as mentioned above. Therefore, we inspected the interfaces with different tilt angles under EM, and found that the zigzag pattern could always be visualized at certain tilt angles for different interfaces (Fig. 2B), suggesting that γB4-ΔIC was stably assembled into an ordered structure at the interfaces.

**Figure 2.**
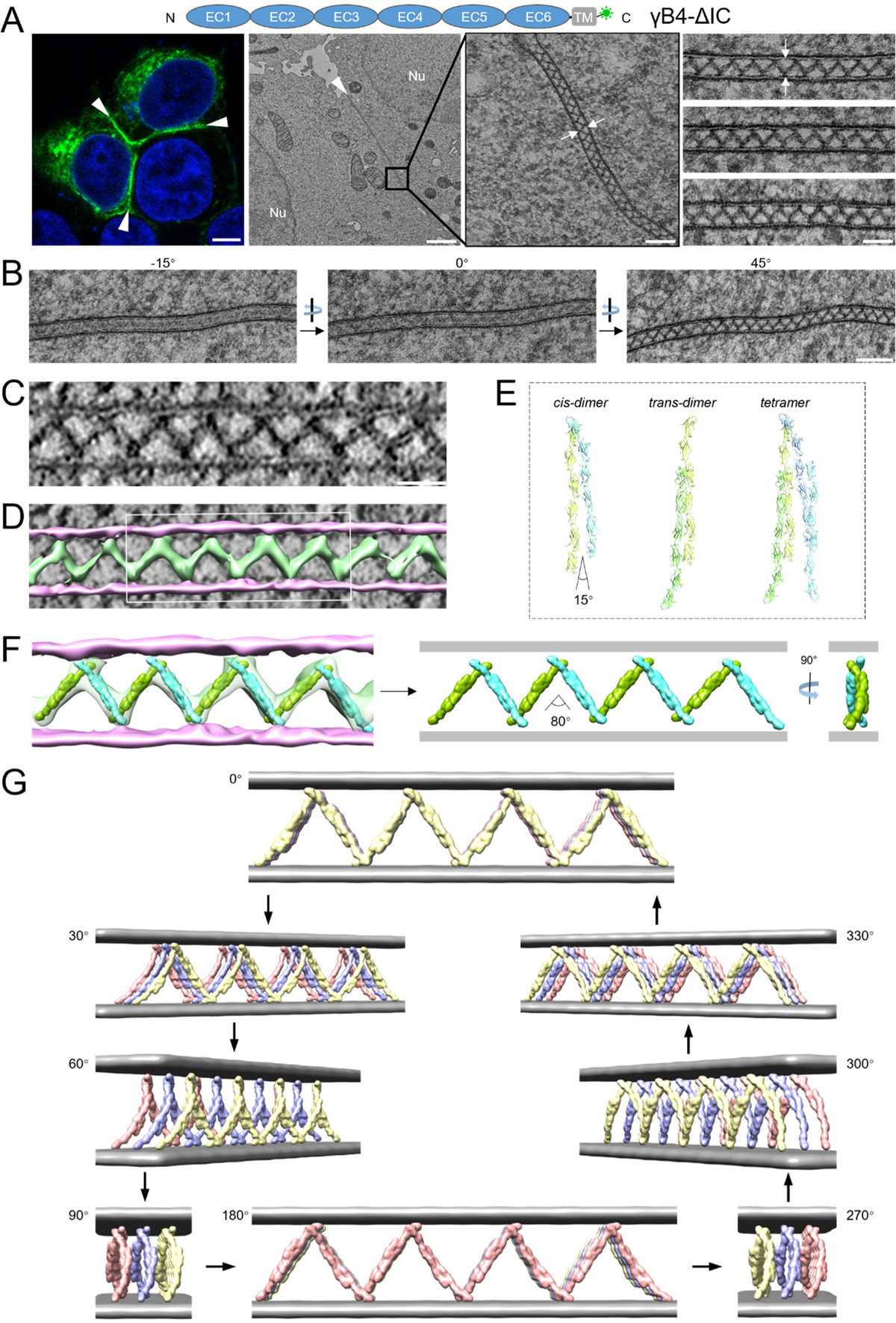
Microscopic images and a tomographic model of the cell adhesion interface by γB4-ΔIC (A) A schematic diagram of the domain arrangement of γB4-ΔIC is shown on the top, the GFP tag is shown in green. A confocal fluorescent image of an adhesion interface (white arrowheads) by γB4-ΔIC is shown on the left (scale bar, 5 μm). EM images of an adhesion interface (white arrowhead) (scale bar, 1 μm) with a zoom-in view (white arrows) (scale bar, 100 nm) are shown in the middle. A gallery of the γB4-ΔIC mediated adhesion interfaces (white arrows) is shown on the right (scale bar, 50 nm). (B) EM images of a γB4-ΔIC mediated adhesion interface visualized at different tilt angles (scale bar, 100 nm). (C) A tomographic slice of a γB4-ΔIC mediated adhesion interface (scale bar, 35 nm). (D) A segmentation model of the tomogram of the γB4-ΔIC mediated adhesion interface shown in (C). The cell membranes and the densities in between are colored in pink and green, respectively. (E) The *trans*- and *cis*-dimers and tetramer of γB4 ectodomain found in crystals. The monomers are colored in light green, green, cyan or blue. (F) The segmentation model shown in (D, white rectangle) is fitted with the *trans*-dimers of the ectodomain of γB4 (green or cyan) (left), revealing the assembly pattern of γB4-ΔIC between cell membranes (right). (G) A 3D tomographic model of the assembly of γB4-ΔIC at the adhesion interfaces. The cell membranes are colored in gray. γB4-ΔIC are shown in yellow, blue or red.

To further characterize the zigzag pattern of γB4-ΔIC at the interfaces, EM tilt series were collected for tomographic reconstruction. In the tomograms, the zigzag pattern could be seen clearly (Fig. 2C), and the intermembrane distance of the adhesion interfaces mediated by γB4-ΔIC was about 28 nm, in contrast to the distance of 34 nm by γB4-FL (Fig.7E). To build an assembly model of γB4-ΔIC, the tomograms were segmented (Fig. 2D), and the crystal structure of the ectodomain of γB4 was docked into the segmented tomograms, revealing the assembly pattern of γB4-ΔIC between cell membranes (Fig. 2E, F). During model fitting, the crystallographic *trans*-dimers of the ectodomain of γB4 matched the tomographic density reasonably well, suggesting that *trans-*dimeric interaction was maintained at the adhesion interfaces. By contrast, the *cis-*dimer in the crystals could not be fitted into the tomograms directly unless the angle between the two monomers of the *cis*-dimer increased from 15 degree to 80 degree (Fig. 2E, F), which would result in a reduction of the intermembrane distance to 28 nm, as observed in the tomograms (Fig. 2C).

A 3D fitting model of γB4-ΔIC was generated according to the tomogram (Fig. 2G, S3 and Mov.1). In the model, the ectodomain of γB4 formed an ordered zigzag pattern between cell membranes which differs from the pattern found in crystal packing (Brasch et al., 2019). The *trans* interaction of γB4 ectodomain is retained, and arrays of γB4-ΔIC were arranged in parallel at the adhesin interfaces (Fig. 2G), which is in agreement with the serial EM sections of the interfaces. The transition from the crystal packing of γB4 ectodomain to the zigzag pattern between cell membranes can be achieved by increasing the angle of the *cis*-dimers like an extendable fence (Fig. 2E, F, G).

### EC5 is important for the zigzag pattern formation of γB4-ΔIC

The zigzag pattern formed by γB4-ΔIC between cell membranes suggests that the ectodomain of γB4 can self-assembled into ordered structure in membrane environment in the absence of IC. Among the EC domains of γB4, EC1-4 are involved in *trans* dimeric interaction, which is retained in the *in situ* assembly of γB4-ΔIC. By contrast, the *cis* dimeric interaction mediated by EC5-6 has changed significantly, implying that they may play a major role in the zigzag pattern formation. Therefore, we made a chimeric molecule where EC5-6 of γB4 are substituted by EC5-6 of γB6 and inspected the pattern formation at the adhesion interfaces (Fig. 3A). The EM data showed that the zigzag pattern disappeared when EC5-6 of γB4 were replaced, confirming the importance of EC5-6 in the assembly (Fig. 3A). Then we generated two chimeric molecules, where either EC5 or EC6 of γB4 was substituted, the resulting images showed that the substitution of EC5 disrupted the zigzag pattern formation (Fig. 3B), whereas the substitution of EC6 had no impact on the pattern formation (Fig. 3C), suggesting that EC5 is crucial for the pattern formation of γB4-ΔIC.

**Figure 3.**
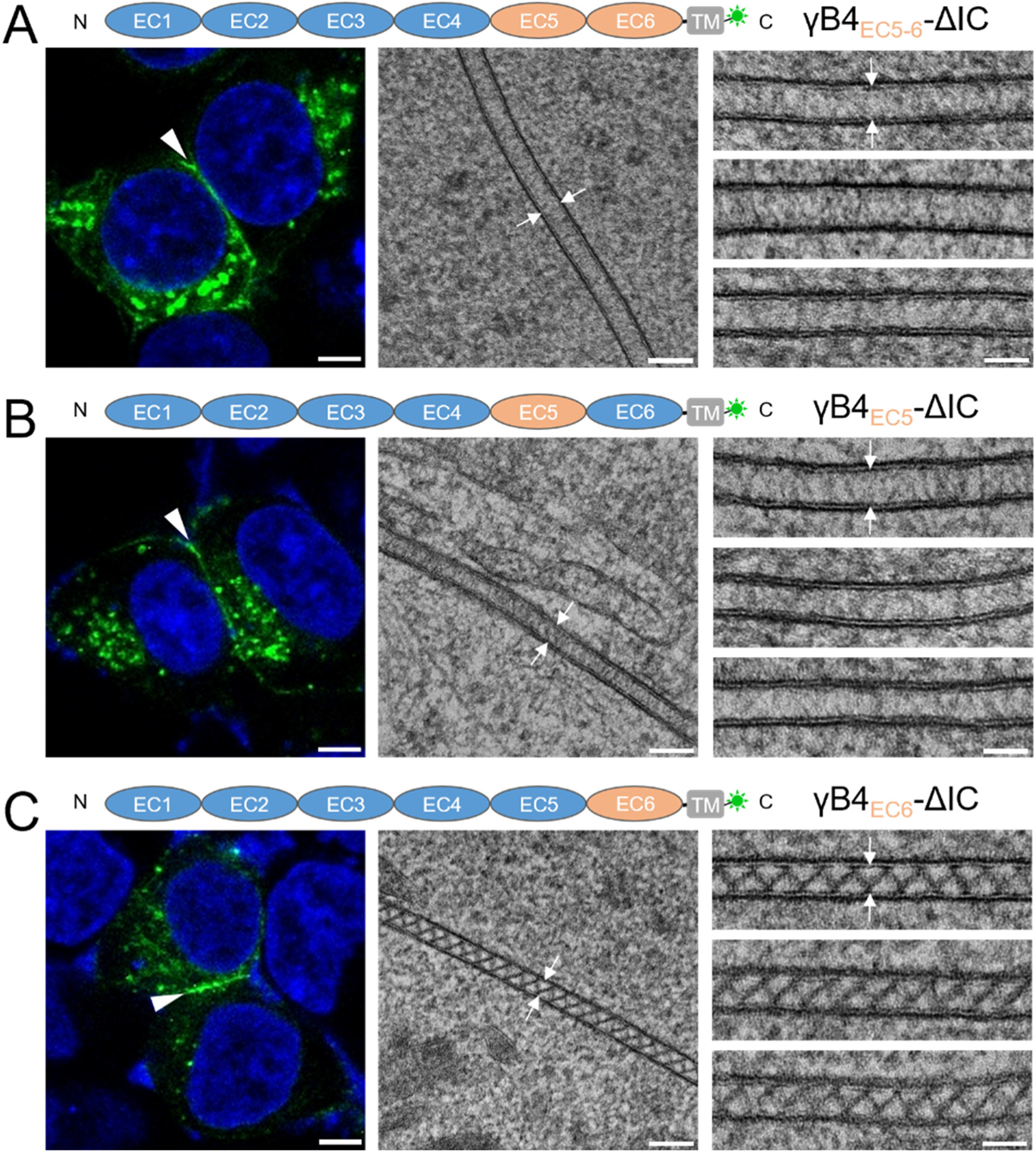
Microscopic images of the cell adhesion interfaces by the substitutional mutants of γB4-ΔIC (A) A schematic diagram of a substitutional mutant of γB4EC5-6-ΔIC is shown on the top. A confocal fluorescent image of an adhesion interface (white arrowhead) by the mutant is shown on the left. An EM image of an adhesion interface (white arrows) is shown in the middle. A gallery of the γB4EC5-6-ΔIC mediated adhesion interfaces (white arrows) is shown on the right. (B) A schematic diagram of a substitutional mutant of γB4EC5-ΔIC is shown on the top. A confocal fluorescent image of an adhesion interface (white arrowhead) by the mutant is shown on the left. An EM image of an adhesion interface (white arrows) is shown in the middle. A gallery of the γB4EC5-ΔIC mediated adhesion interfaces (white arrows) is shown on the right. (C) A schematic diagram of a substitutional mutant of γB4EC6-ΔIC is shown on the top. A confocal fluorescent image of an adhesion interface (white arrowhead) by the mutant is shown on the left. An EM image of an adhesion interface (white arrows) is shown in the middle. A gallery of the γB4EC6-ΔIC mediated adhesion interfaces (white arrows) is shown on the right. Scale bar, 5 μm (left), 100 nm (middle), 50 nm (right).

To identify the residues on EC5 that might be important for the assembly, we did a sequence alignment of EC5 between γB4 and γB6, and the result showed a high sequence identity (87%) except in four regions: T451-V453, Q484-Y488, E497 and H535-S537 (Fig. 4A). Then we made four mutants of γB4-ΔIC by replacing the corresponding residues, including T451Q/V453S, Q484H/Y488S, E497K and H535Q/S537K (Fig. 4A). The mutants were applied for EM visualization of the adhesion interfaces, and the results showed that all the mutants except E497K, retained the zigzag pattern at the cell interfaces (Fig. 4B-E), suggesting that E497 of EC5 might play an important role in the ordered assembly of γB4-ΔIC.

**Figure 4.**
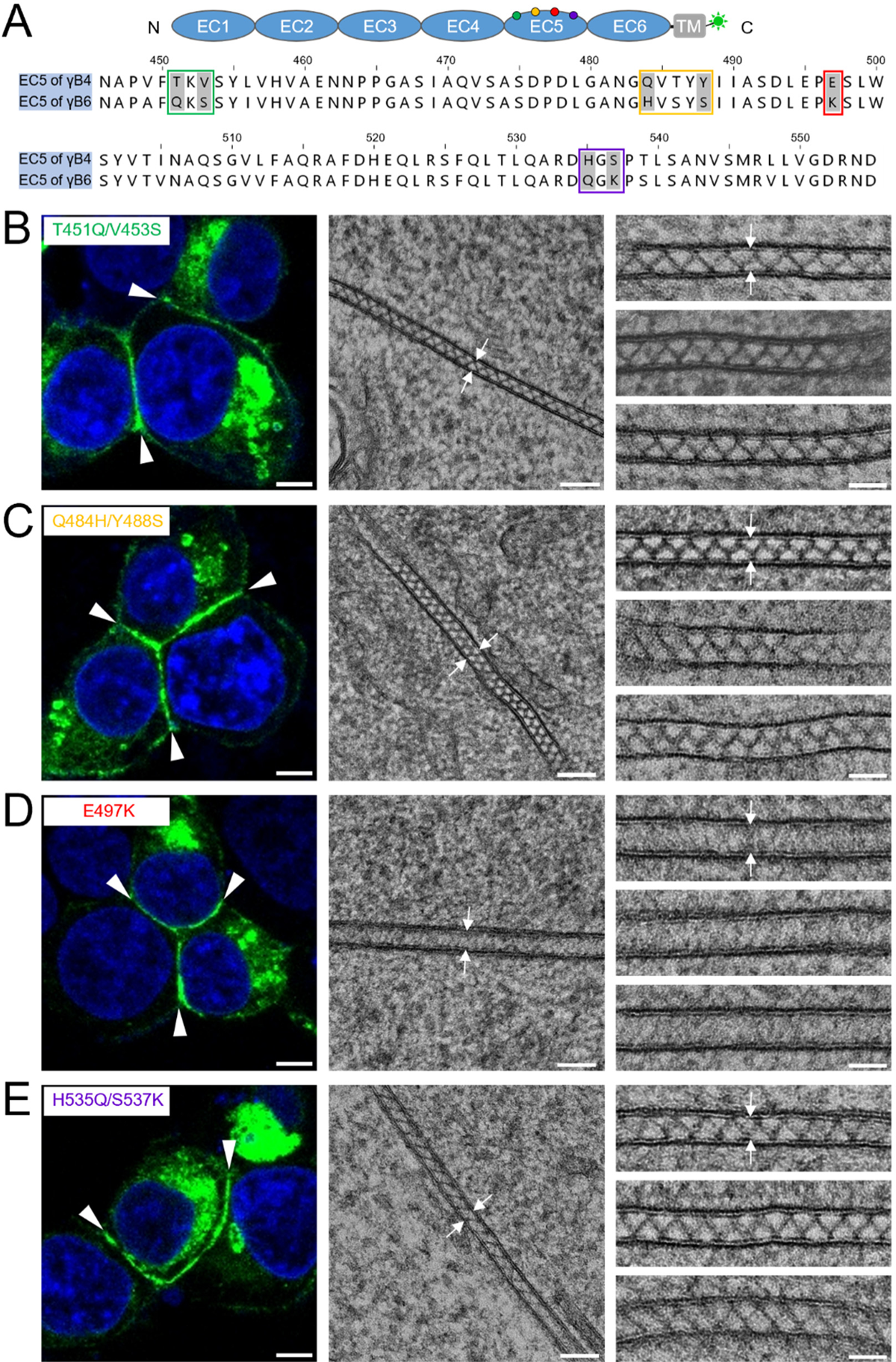
Microscopic images of the cell adhesion interfaces by the EC5 mutants of γB4-ΔIC (A) A schematic diagram of γB4-ΔIC is shown on the top. The sequence alignment of EC5 from γB4 and γB6 is shown at the bottom. The residue differences between γB4 and γB6 on EC5 are labeled in green, yellow, red or purple. (B) A confocal fluorescent image of an adhesion interface (white arrowheads) by the mutant T451Q/V453S is shown on the left. An EM image of an adhesion interface (white arrows) is shown in the middle. A gallery of the mutant mediated adhesion interfaces (white arrows) is shown on the right. (C) A confocal fluorescent image of an adhesion interface (white arrowheads) by the mutant Q484H/Y488S is shown on the left. An EM image of an adhesion interface (white arrows) is shown in the middle. A gallery of the mutant mediated adhesion interfaces (white arrows) is shown on the right. (D) A confocal fluorescent image of an adhesion interface (white arrowheads) by the mutant E497K is shown on the left. An EM image of an adhesion interface (white arrows) is shown in the middle. A gallery of the mutant mediated adhesion interfaces (white arrows) is shown on the right. (E) A confocal fluorescent image of an adhesion interface (white arrowheads) by the mutant H535Q/S537K is shown on the left. An EM image of an adhesion interface (white arrows) is shown in the middle. A gallery of the mutant mediated adhesion interfaces (white arrows) is shown on the right. Scale bar, 5 μm (left), 100 nm (middle), 50 nm (right).

### Cis-interaction of the in situ assembly of γB4-ΔIC

Tomographic model fitting suggested that the angle of the crystallographic *cis*-dimer increased from 15 degree to 80 degree (Fig. 5A, B). In the crystal structure, E497 locates on the surface of EC5 (Fig. 5A). Following the angle change of the *cis*-dimer, E497 might be able to approach a positively charged region on the surface of EC6 from the other monomer, which may provide electrostatic interaction to stabilize the zigzag pattern and could also explain the disruption of the zigzag pattern by the single mutation E497K (Fig. 5B).

**Figure 5.**
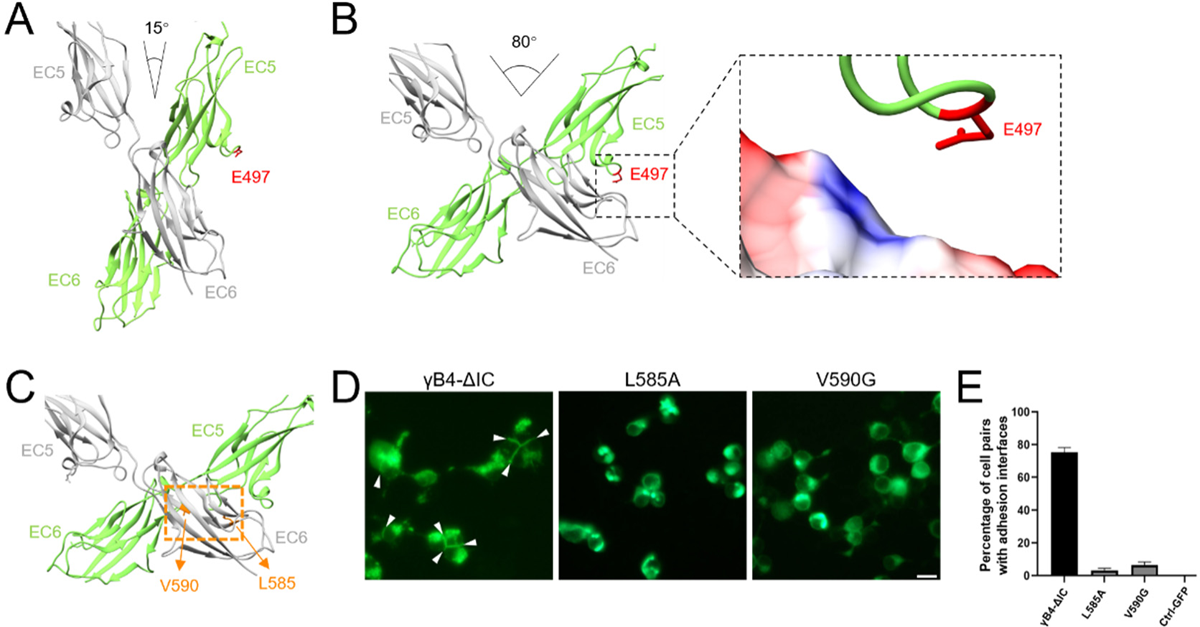
*Cis*-dimeric interaction of the *in situ* assembly of γB4-ΔIC (A) *Cis*-dimeric interaction of γB4 ectodomain in the crystals. EC5 and EC6 from the two monomers are colored in gray or green. The position of E497 from one of the monomers is labeled. (B) *Cis*-dimeric interaction of γB4-ΔIC on the cell surface. EC5 and EC6 from the two monomers are colored in gray or green. The position of E497 from one of the monomers is labeled. A positively charged region (blue) from EC6 that may be approached by E497 during the *in situ* assembly of γB4-ΔIC is also shown (dashed rectangles). (C) The potential *cis*-dimeric interface of γB4-ΔIC on the cell surface (dashed orange rectangle). The positions of L585 and V590 are labeled. (D) The fluorescent images of cell adhesion mediated by γB4-ΔIC and γB4 mutants (L585A and V590G). The adhesion interfaces are indicated by white arrowheads (scale bar, 15 μm). (E) The statistics of the adhesion interfaces by γB4-ΔIC and γB4 mutants (L585A and V590G). The GFP transfected cells are applied as a control.

According to the published data, residues L585 and V590 locate at the *cis-* dimeric interface of γB4 ectodomain in the crystal structure and are important for forming cell adhesion (Goodman et al., 2017; Goodman et al., 2016b). Here we also made mutants of the two residues, L585A and V590G, on γB4-ΔIC (Fig. 5C), and indeed, no adhesion interface was identified for the cells transfected with these two mutants by fluorescent microscopy (Fig. 5D, E), similar to the previous observation (Goodman et al., 2017; Goodman et al., 2016b), implying that these residues might also be important for the assembly of γB4-ΔIC on the cell surface. Taken together, it appears that the *cis*-dimeric interface found in the crystal structure may act as a “hinge” maintained by hydrophobic interactions (Fig. 5C), while E497 may interact with the neighboring monomers through charge interaction and stabilize the large opening angle of the *cis-*dimers in the zigzag pattern of γB4-ΔIC at the adhesion interfaces (Fig. 5B)

### Intracellular domain regulates the in situ assembly of γB4

As shown above, the *in situ* assembly patterns of γB4-FL and γB4-ΔIC are significantly different, suggesting that the intracellular domain of γB4 is involved in regulating the organization of γB4 at the adhesion interfaces. In fact, the published data showed that the intracellular domains of cPcdhs could interact with each other (Murata et al., 2004; O’Leary et al., 2011; Shonubi et al., 2015), therefore may affect the assembly of the ectodomains of cPcdhs. To verify the interaction between the intracellular domains on the cell membrane, we co-transfected cells with γB4-FL (fused with RFP) and IC of γB4 (including TM and IC, fused with GFP), confocal images showed that IC could co-localize with γB4-FL at the interfaces (Fig. 6A).

**Figure 6.**
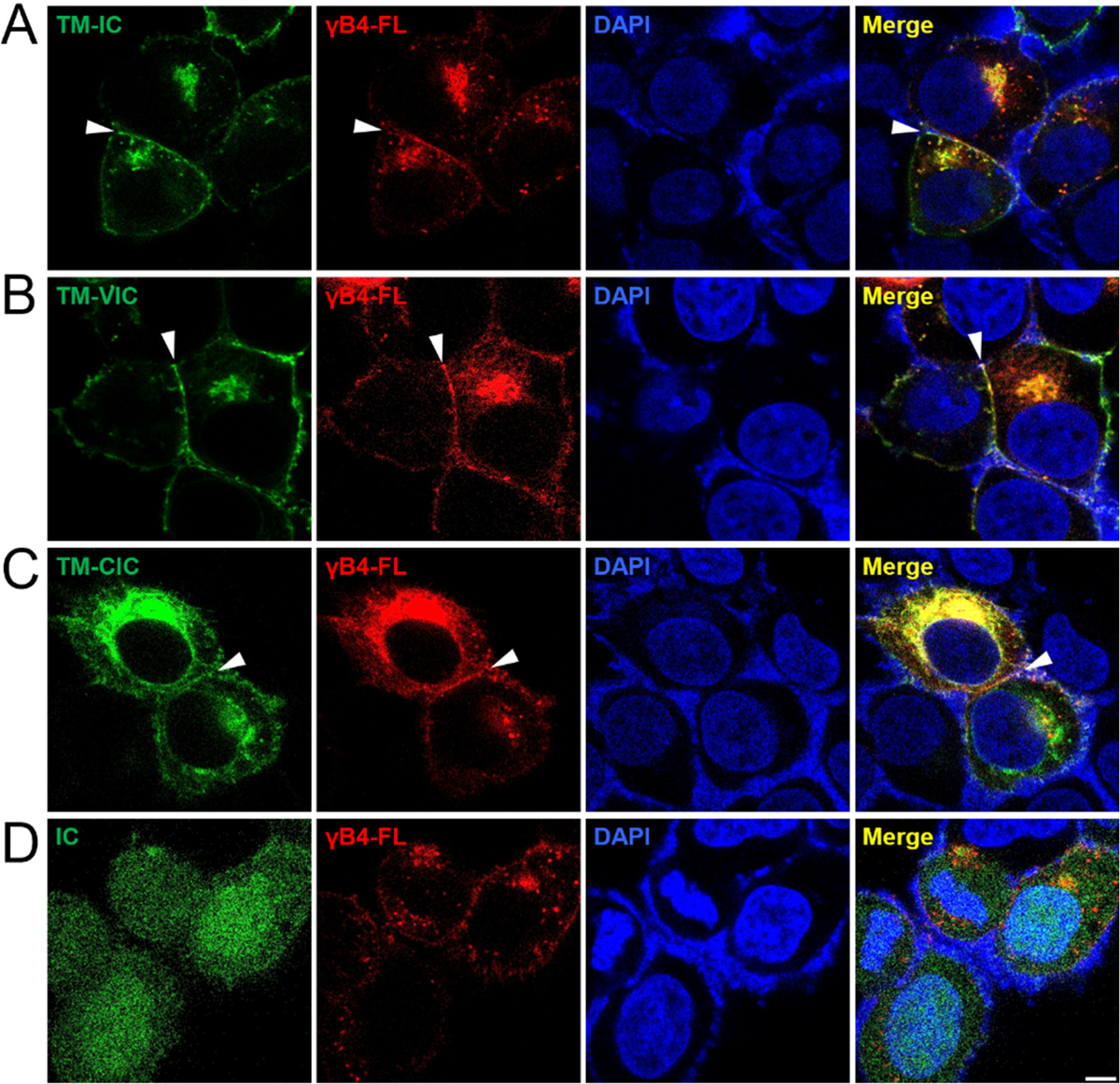
Confocal fluorescent images of the cells co-transfected with γB4-FL and the IC mutants of γB4 (A) Confocal images of the cells co-transfected with γB4-FL (red) and TM-IC (green). (B) Confocal images of the cells co-transfected with γB4-FL (red) and TM-VIC (green). (C) Confocal images of the cells co-transfected with γB4-FL (red) and TM-CIC (green). (D) Confocal images of the cells co-transfected with γB4-FL (red) and IC (green). DAPI is applied to stain the nucleus. The adhesion interfaces are indicated by white arrowheads. Scale bar, 5 μm.

Furthermore, since IC of γB4 contains both VIC and CIC, we co-transfected γB4-FL (fused with RFP) with VIC or CIC (including TM and VIC or CIC, fused with GFP), and both confocal images displayed co-localization of VIC and CIC with γB4-FL (Fig. 6B, C), confirming that the intracellular domain of γB4 could interact with each other on the cell membrane during adhesion. In addition, the confocal images showed that in the absence of TM, IC alone distributed all over the cells including nucleus (Fig. 6D), which was consistent with the published data showing that cleaved fragments of the intracellular domains of cPcdhs had nuclear localization (Bonn et al., 2023; Emond and Jontes, 2008; Haas et al., 2005). These results suggest that the *cis*-interaction between the intracellular domains may not be very strong and only occur locally when they stay on the cell membrane.

To further explore the impact of the intracellular domain on the *in situ* assembly of γB4, we generated two IC-truncation mutants, γB4-ΔCIC and γB4-ΔVIC, where CIC or VIC was removed from IC. Fluorescent microscopy showed that both mutants could mediate adhesion of the transfected cells (Fig. 7A, C). EM images and the tomograms showed that the two mutants did not form ordered assemblies between cell membranes (Fig. 7A-D), and the intermembrane distances of the two mutants determined by EM were about 33 nm, similar to that of γB4-FL (Fig. 7E). However, the intermembrane tomographic densities of the two mutants were different, it appeared that more molecules were recruited at the interfaces than γB4-FL, and γB4-ΔVIC seemed to have more molecules at the interface than γB4-ΔCIC (Fig. 1C, 7B, 7D, S4). This is in agreement with the previous data showing that the intracellular domains may restrict accumulation of cPcdhs at cell-cell contacts (Fernandez-Monreal et al., 2009; Schreiner and Weiner, 2010), and suggests that partial deletion of the intracellular domain may reduce its impact on the organization of the ectodomain, and VIC may have larger impacts on the assembly than CIC, which is not surprising as VIC locates closer to the cell membrane than CIC. In addition, we also evaluated the efficiency of adhesion formation mediated by γB4-FL and the IC-truncation mutants. The resulting statistics showed that γB4-ΔIC had the highest adhesion efficiency, implying that the zigzag pattern was preferred for the ectodomain, and the intracellular domain reduced adhesion efficiency significantly (Fig. 7F). Deletion of VIC or CIC increased the adhesion formation and may also partially recover the assembly of the ectodomain at the interfaces (Fig. 7F, S4). In addition, structural prediction by AlphaFold (Jumper et al., 2021; Varadi et al., 2021) shows that the intracellular domains of cPcdhs are rather flexible without secondary structure, how they regulate the molecular organization *in situ* still need further investigation in the future.

**Figure 7.**
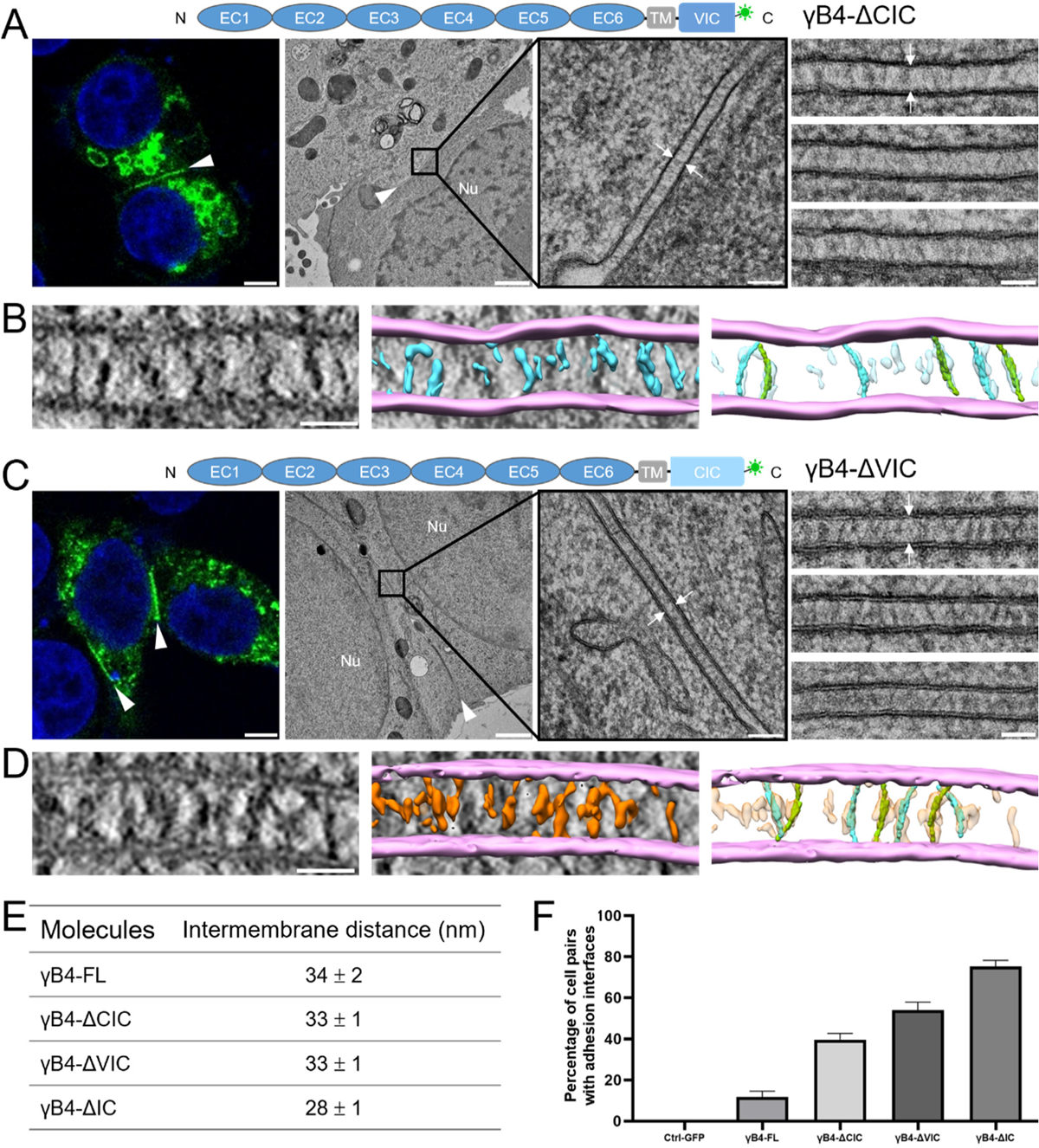
Microscopic images and the statistics of the adhesion interfaces by the IC-truncation mutants of γB4. (A) A schematic diagram of γB4-ΔCIC is shown on the top. A confocal fluorescent image of an adhesion interface (white arrowhead) by γB4-ΔCIC is shown on the left (scale bar, 5 μm). EM images of an adhesion interface (white arrowhead) (scale bar, 1 μm) with a zoom-in view (white arrows) (scale bar, 100 nm) are shown in the middle. A gallery of the γB4-ΔCIC mediated adhesion interfaces (white arrows) is shown on the right (scale bar, 50 nm). (B) A tomographic slice of a γB4-ΔCIC mediated adhesion interface (left) (scale bar, 35 nm) and a segmentation model of the tomogram (middle). The cell membranes and the densities in between are colored in pink and cyan, respectively. The densities are tentatively docked with the *trans*-dimers of the ectodomain of γB4 (green or cyan) (right). (C) A schematic diagram of γB4-ΔVIC is shown on the top. A confocal fluorescent image of an adhesion interface (white arrowhead) by γB4-ΔVIC is shown on the left (scale bar, 5 μm). EM images of an adhesion interface (white arrowhead) (scale bar, 1 μm) with a zoom-in view (white arrows) (scale bar, 100 nm) are shown in the middle. A gallery of the γB4-ΔVIC mediated adhesion interfaces (white arrows) is shown on the right (scale bar, 50 nm). (D) A tomographic slice of a γB4-ΔVIC mediated adhesion interface (left) (scale bar, 35 nm) and a segmentation model of the tomogram (middle). The cell membranes and the densities in between are colored in pink and orange, respectively. The densities are tentatively docked with the *trans*-dimers of the ectodomain of γB4 (green or cyan) (right). (E) Statistics of the intermembrane distances of the *in situ* assemblies of γB4-FL, γB4-ΔCIC, γB4-ΔVIC and γB4-ΔIC. (F) The statistics of the adhesion interfaces by γB4-FL, γB4-ΔCIC, γB4-ΔVIC and γB4-ΔIC. The GFP transfected cells are applied as a control. The data for γB4-ΔIC is also shown in Fig. 5E.

## Discussion

Cell adhesion molecules are important for mediating cell-cell contacts, their structures, especially the ectodomains, have been studied extensively by X-ray crystallography in the past decades (Honig and Shapiro, 2020). Recent developments in EM provide opportunities to visualize their *in situ* organizations on the cell surface (Chang et al., 2018; McCafferty et al., 2024), which advances the understanding of the mechanisms of cell adhesion. The *in situ* studies of IgSF adhesion molecules showed that their assemblies are mainly regulated by the ectodomains through *trans* and *cis* as well as membrane interactions, and some molecules may form ordered assemblies at the interfaces (Guo et al., 2021; Tang et al., 2018). It has been proposed that the ordered assembly of the ectodomains may affect the downstream signaling or cytoskeletal organization (Boni et al., 2022; Honig and Shapiro, 2020). But whether ordered organization is a general feature for the assemblies of adhesion molecules still needs further investigation.

The crystal structure of the ectodomain of γB4 shows that EC1-4 mediate *trans* dimer formation (Goodman et al., 2022; Goodman et al., 2016a; Goodman et al., 2016b) and EC5-6 are responsible for forming asymmetric *cis* dimers, where EC5-6 of a monomer interacts with EC6 of the other monomer (Goodman et al., 2022; Goodman et al., 2017). The alternate *trans* and *cis* dimerizations can produce a zipper-like assembly in crystal packing (Fig. 8A), and a similar pattern has been observed in a liposome model for γB6 (Brasch et al., 2019). However, the *in situ* imaging of the adhesion interfaces by γB4-FL does not show an ordered assembly pattern, suggesting that *in situ* assembly could be more complex and some interactions might be missing in crystal packing. Indeed, when the intracellular domain of γB4 is removed, its intermembrane organization changes dramatically by forming an ordered zigzag pattern. The tomographic model shows that the zigzag pattern can be generated by the *trans* and *cis* dimers of the ectodomain of γB4, where the *trans* dimer is similar to that in the crystals, but the *cis* dimer adopts a larger opening angle between the monomers, suggesting that the ectodomain of γB4 can self-assemble into ordered structures on the cell surface, which might be driven by the forces not identified in crystals. The subsequent mutagenesis data show that the *cis* dimeric interface formed between EC5-6 and EC6 is probably or at least partially retained, and E497 on EC5 may facilitate the opening of the *cis*-dimers through charge interaction and stabilize the zigzag pattern.

**Figure 8.**
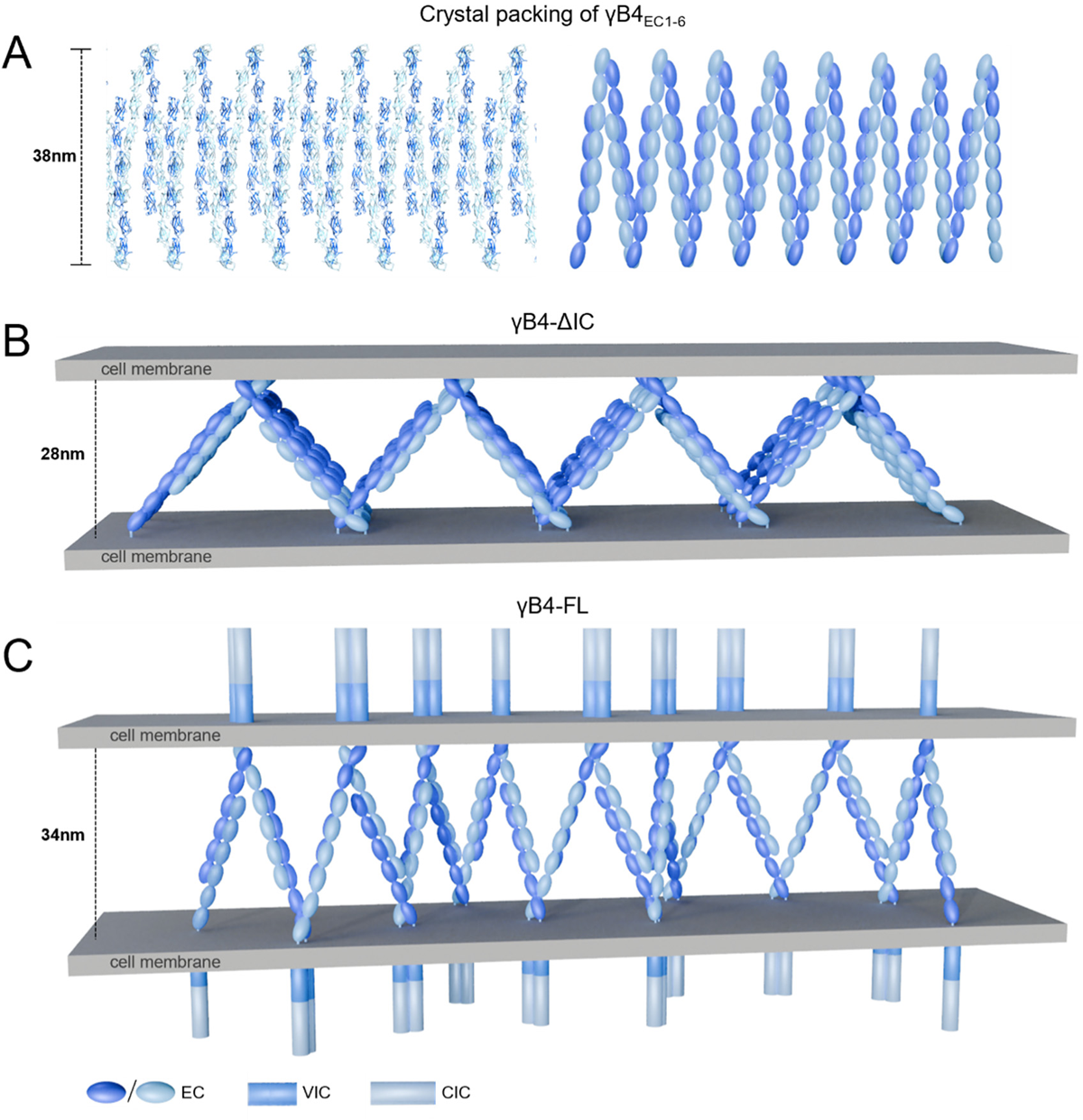
Crystal packing of the ectodomain of γB4 and the *in situ* assembly models for γB4-ΔIC and γB4-FL (A) Crystal packing of the ectodomain of γB4 (left). A schematic model is shown on the right. (B) An *in situ* assembly model of γB4-ΔIC. (C) An *in situ* assembly model of γB4-FL.

The different assembly patterns of γB4-FL and γB4-ΔIC suggest that the intracellular domain also regulates the organization of γB4 between cell membranes, which may not be entirely unexpected as previous data have shown that the intracellular domains of cPcdhs can interact with each other (Murata et al., 2004; Shonubi et al., 2015) and is also in agreement with our fluorescent observation. Moreover, tomographic data suggest that both VIC and CIC could affect the assembly of γB4 on the cell surface, probably mainly on the *cis* organization of the molecules, and VIC seems to have larger impacts on the assembly due to its proximal location to the membrane. Based on these information, the schematic models for the *in situ* assembly of γB4-ΔIC and γB4-FL are generated (Fig. 8B, C).

Published data have shown that *cis*-interactions between the ectodomains of cPcdhs are important for homophilic combinatorial cell recognition in mediating self-avoidance (Brasch et al., 2019; Rubinstein et al., 2017; Rubinstein et al., 2015; Wiseglass et al., 2023). Here we show that the intracellular domain of γB4 also regulates the intermembrane assembly pattern, especially the *cis* organization of the molecule, suggesting that it might be crucial in establishing homophilic adhesion between cells. Although the *in situ* assembly model of γB4-FL represents a simple case with only γB4 on the cell surface (Fig. 8C), it may provide clues for the situation where combinatorial expression of cPcdhs occurs on the membrane. In fact, the sequences of the intracellular domains of cPcdhs are rather diverse, all isoforms have VIC, which contains variable sequences, and α- and γ-cPcdhs have cluster-specific CIC, which is missing in β-cPcdhs (Wu and Maniatis, 1999; Wu et al., 2001).

Evidence has also shown that the intracellular domains of different cPcdhs could interact with each other (Murata et al., 2004; Shonubi et al., 2015). Therefore, it might be possible that the *cis*-interactions among the isoforms may help to generate complex but specific assembly patterns like 2D barcodes on the cell surfaces for each set of cPcdhs and lead to homophilic adhesion between cells, and a single isoform mismatch would result in a different assembly pattern and disrupt the adhesion for self-avoidance. Overall, these data suggest that both ectodomains and intracellular domains of cPcdhs contribute to the homophilic adhesion between cells, but their exact roles and mechanisms still need further investigation in the future.

## Material and Methods

### Preparation of DNA constructs

cDNA of the full-length mouse cPcdh-γB4 (residues 1-912, residue number includes the signal peptide) was cloned into pCMV expression vector fused with a GFP or mCherry tag at the C-terminus. The truncation mutants of IC (residues 721-912), TM-IC (residues 1-30, 663-912), TM-VIC (residues 1-30, 663-788), TM-CIC (residues 1-30, 663-720, 789-912), γB4-ΔIC (residues 1-720), γB4-ΔVIC (residues 1-720, 789-912), γB4-ΔCIC (residues 1-788) were also subcloned into pCMV vector similarly. For the domain substitution mutants, EC5(residues 446-555) and/or EC6 (residues 556-662) of γB4 were replaced by the EC5 (residues 448-557) and/or EC6 (residues 558-664) of γB6 by homologous recombination using the *ClonExpress MultiS One Step Cloning kit* (Vazyme, C113-01). The single or double mutants of γB4-ΔIC, including T451Q/V453S, Q484H/Y488S, E497K, H535Q/S537K, L585A and V590G, were constructed by PCR.

### Confocal microscopy

HEK293T cells were cultured on coverslips coated with Poly-L-lysine (Sigma, P4707-50ML). Plasmid constructs were transfected into the cells by using Lipofectamine 2000 reagent (Invitrogen, 11668019). After 24 hours, transfected cells were fixed with 4% paraformaldehyde, permeabilized with 0.5% Triton-X and mounted with antifade mounting medium with DAPI (Beyotime, P0131-25ml).

Images were acquired on a confocal microscope Leica TCS SP8. For the adhesion formation statistics, cells were cultured on multiple well plates and transfected with the γB4 constructs. After 24 hours, cells were visualized under fluorescence microscope ECLIPSE Ts2, the percentage of highlighted fluorescent interfaces appeared in the pairs of neighboring cells were counted. The experiments were repeated four times (five visional fields each time). The data were analyzed by GraphPad Prism 9.0 and plotted as mean ± SE.

### EM sample preparation

Sapphire discs were marked by carbon evaporation and coated with poly-L-lysine for cell culture. HEK293T cells were transfected with the target constructs. After 24 hours, the sapphire discs were transferred to specimen holders and covered by aluminum planchettes with 25-μm inner depth and filled with hexadecane. The specimens were then loaded on a Wohlwend HPF Compact 2 high-pressure freezer (M.Wohlwend GmbH) for HPF. Frozen specimens were transferred into cryotubes containing 0.1% uranyl acetate, 0.6% water, and 1% osmium tetroxide in acetone at liquid nitrogen temperature. Freeze substitutions were completed as previously described (Chang et al., 2018; Guo et al., 2021; Tang et al., 2018), cells were then embedded into resin blocks and solidified. The resin blocks were subjected to thin sectioning on a Leica EM UC7 ultra-microtome. Ultrathin sections of 100 nm thickness were collected onto formvar-coated copper grids with an evaporated carbon film and stained with 3% uranyl acetate at 4℃ for 7 minutes, then by lead citrate at room temperature for 3 minutes. The stained sections were loaded on a 120 kV Tecnai T12 microscope (Thermo Fisher Scientific) for imaging.

### Electron tomography

Ultrathin sections were loaded onto a FEI Tecnai G2 electron microscope (120 kV) for collecting tomographic tilt series. Single axis tilt series were collected ranging from −60° to 60° with 1.5° increments at a pixel size of 1.71 Å using Xplore3D (FEI). Tomograms were reconstructed using EMAN2.9 and the final tomograms were binned with a resulting pixel size of 6.86 Å (Chen et al., 2019). Fiji (Schindelin et al., 2012) was applied for measuring the intermembrane distances and calculating the histograms of the intermembrane densities in the tomograms. Segmentation was done semi-automatically by EMAN2.9 combined with IMOD (Chen et al., 2017a; Danita et al., 2022).

### Model building

The atomic models were built by docking the crystal structure of the ectodomain γB4 into the segmented tomogram maps manually using UCSF chimera (Pettersen et al., 2004). The *trans*-dimers of γB4 were retained as in the crystal structure and fitted into the tomographic density volumes. The *cis*-dimers were built according to the tomographic density using UCSF chimera (Pettersen et al., 2004). The movie was made by UCSF chimera. The schematic models of γB4-ΔIC and γB4-FL were built using Blender (https://www.blender.org).

## Acknowledgements

We thank the Electron Microscopy and Integrated Laser Microscopy Systems at the National Facility for Protein Science in Shanghai (NFPS), Shanghai Advanced Research Institute, Chinese Academy of Sciences, China, for technical support. This work is supported by National Natural Science Foundation of China (No. 32241022 and 32271243) to Y.H. and we also thank the support from Innovative research team of high-level local universities in Shanghai (SHSMU-ZLCX20212601).

## Supplementary figures

**Figure S1.**
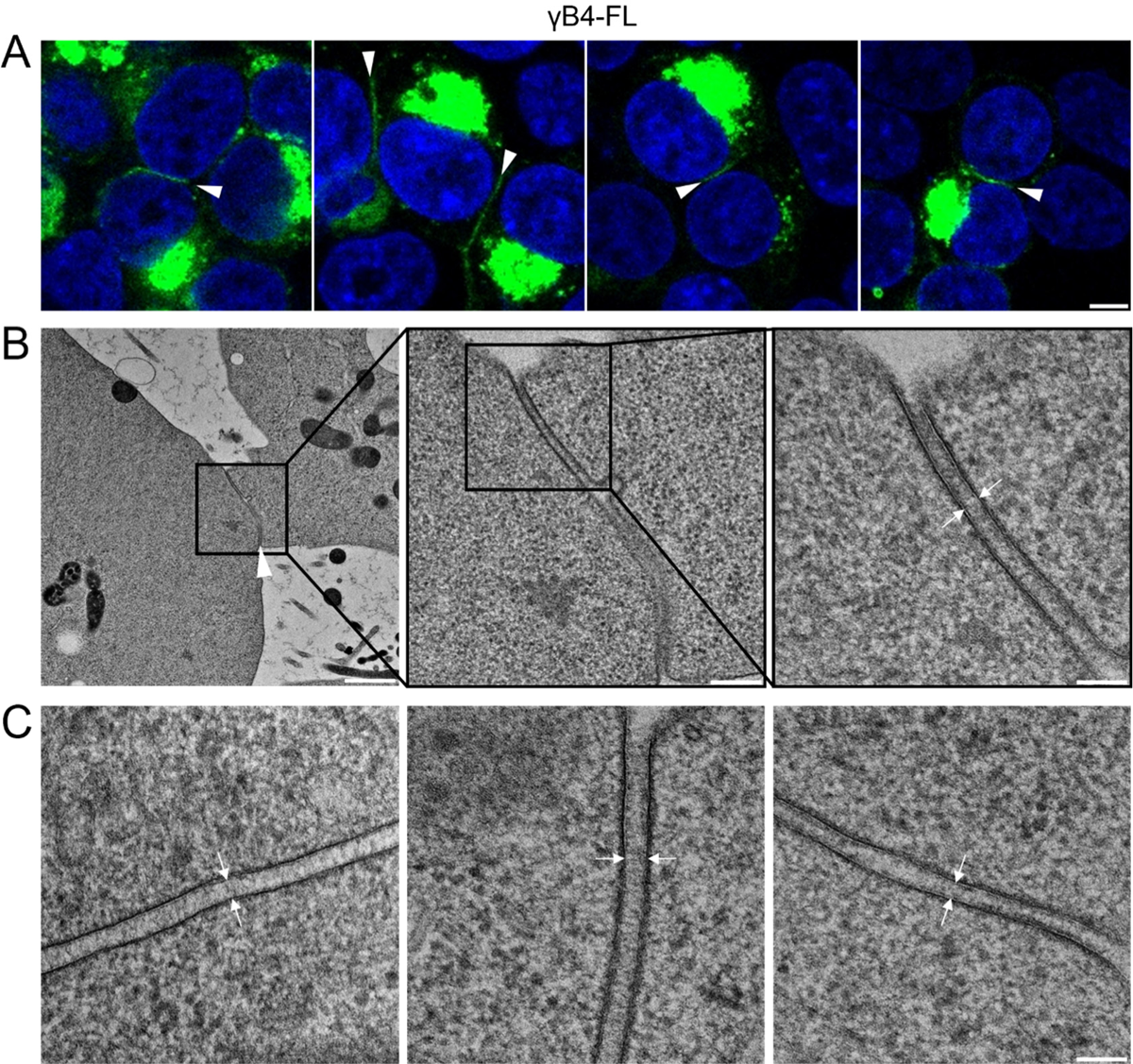
Microscopic images of the cell adhesion interfaces by γB4-FL (A) Confocal fluorescent images of adhesion interfaces (white arrowheads) by γB4-FL Scale bar, 5 μm. (B) EM images of an adhesion interface (white arrowhead, left) by γB4-FL and the zoom-in views (middle and right). Scale bar, 1 μm (left); 250 nm (middle); 100 nm (right). (C) A gallery of the γB4-FL mediated adhesion interfaces (white arrows). Scale bar, 100 nm.

**Figure S2.**
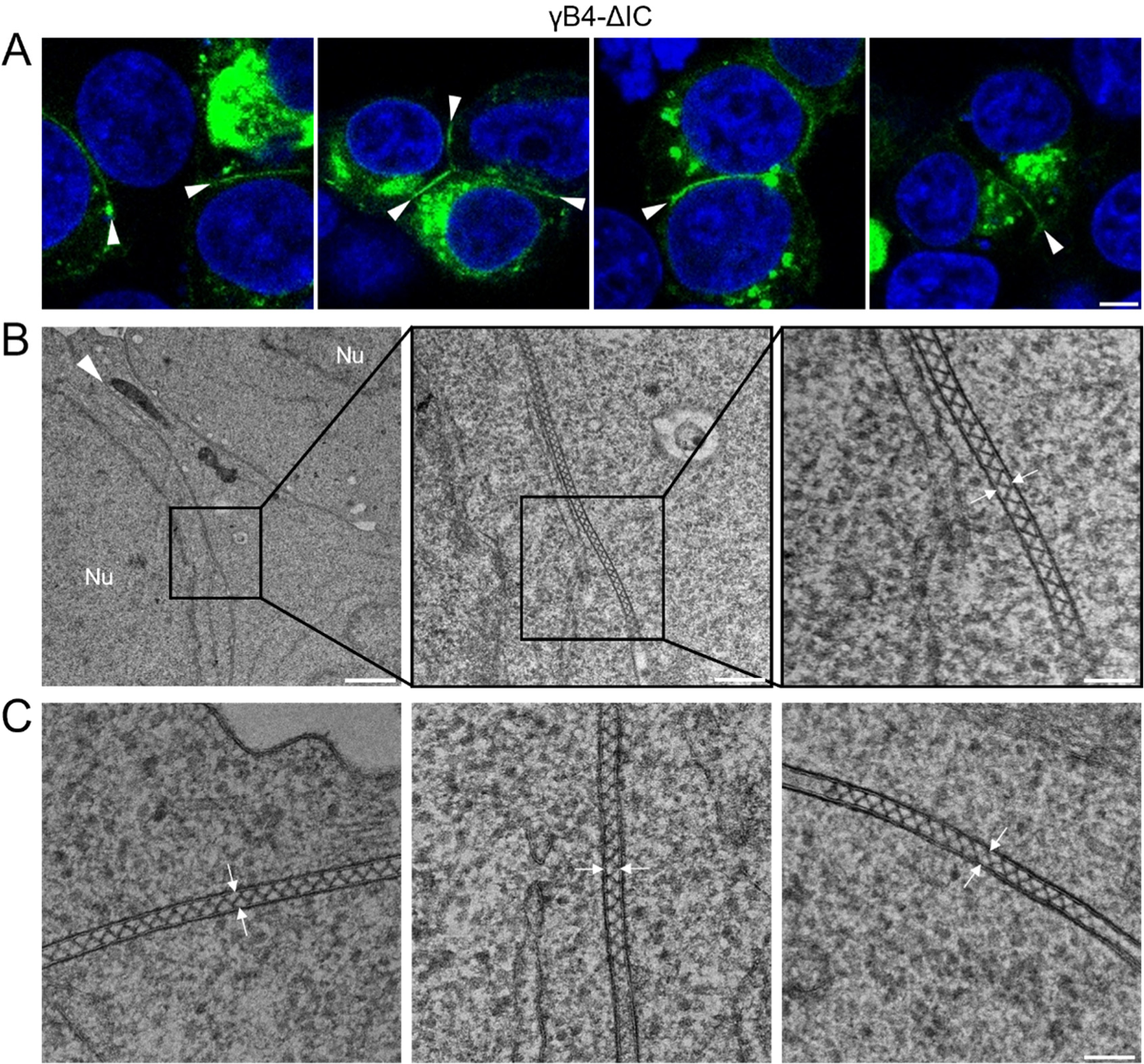
Microscopic images of the cell adhesion interfaces by γB4-ΔIC (A) Confocal fluorescent images of adhesion interfaces (white arrowheads) by γB4-ΔIC. Scale bar, 5 μm. (B) EM images of an adhesion interface (white arrowhead, left) by γB4-ΔIC and the zoom-in views (middle and right). Scale bar, 1 μm (left); 250 nm (middle); 100 nm (right). (C) A gallery of the γB4-ΔIC mediated adhesion interfaces (white arrows). Scale bar, 100 nm.

**Figure S3.**
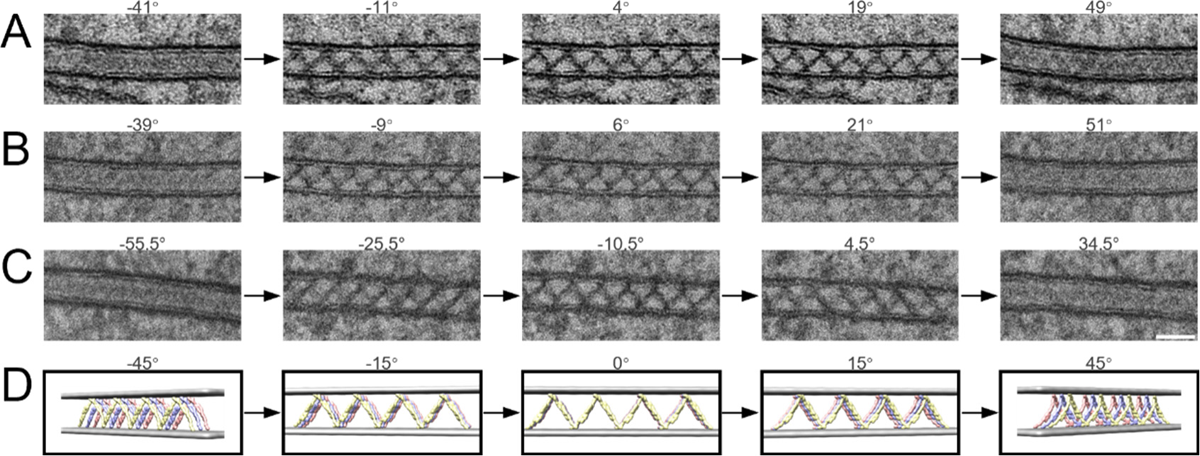
Tomographic tilt series of the cell adhesion interfaces by γB4-ΔIC. (A-C) Three tomographic tilt series of the cell adhesion interfaces by γB4-ΔIC visualized at different tilt angles. Scale bar, 50 nm. (D) The 3D assembly model of γB4-ΔIC visualized at the corresponding tilt angles shown in (A-C).

**Figure S4.**
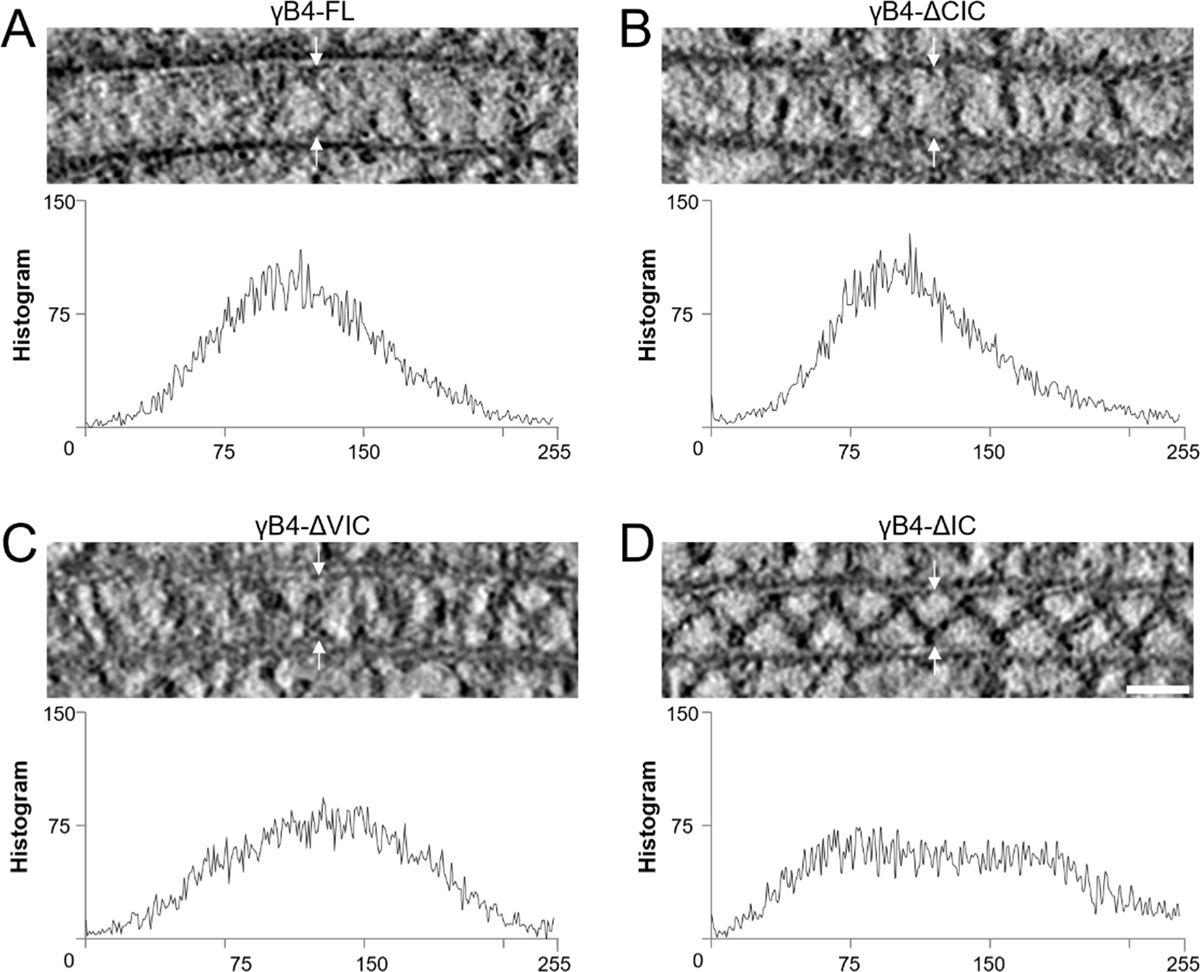
Histograms of the intermembrane tomographic densities of the interfaces by γB4-FL and the IC-truncation mutants of γB4. (A) A tomographic slice of the γB4-FL mediated interface (top, also shown in Fig. 1C) and the corresponding histogram (bottom). (B) A tomographic slice of the γB4-ΔCIC mediated interface (top, also shown in Fig. 7B) and the corresponding histogram (bottom). (C) A tomographic slice of the γB4-ΔVIC mediated interface (top, also shown in Fig. 7D) and the corresponding histogram (bottom). (D) A tomographic slice of the γB4-ΔIC mediated interface (top, also shown in Fig. 2C) and the corresponding histogram (bottom).Scale bar, 35 nm.

**Mov.1** A tomogram and 3D model of the cell adhesion interface by γB4-ΔIC

